# Neurochemical correlates of scene processing in the precuneus/posterior cingulate cortex: a multimodal fMRI and ^1^H-MRS study

**DOI:** 10.1101/422758

**Authors:** Alison G. Costigan, Katja Umla-Runge, C. John Evans, Carl J. Hodgetts, Andrew D. Lawrence, Kim S. Graham

## Abstract

Precuneus/posterior cingulate cortex (PCu/PCC) are key components of a midline network, activated during rest but also in tasks that involve construction of scene or situation models. Despite growing interest in PCu/PCC functional alterations in disease, the underlying neurochemical modulators of PCu/PCC’s task-induced activity are largely unstudied. Here, a multimodal imaging approach was applied to investigate whether inter-individual differences in PCu/PCC fMRI activity, elicited during perceptual discrimination of scene stimuli, were correlated with local brain metabolite levels, measured during resting-state ^1^H-MRS. Forty healthy young adult participants (12 male) completed an fMRI perceptual odd-one-out task for scenes, objects and faces. ^1^H-MRS metabolites N-acetyl-aspartate (tNAA), glutamate (Glx) and γ-amino-butyric acid (GABA+) were quantified via PRESS and MEGA-PRESS scans in a PCu/PCC voxel and an occipital (OCC) control voxel. Whole brain fMRI revealed a cluster in right dorsal PCu/PCC that showed a greater BOLD response to scenes versus faces and objects. When extracted from an independently defined PCu/PCC region of interest, scene activity (versus faces and objects and also versus baseline) was positively correlated with PCu/PCC, but not OCC, tNAA. A complementary fMRI analysis restricted to the PCu/PCC MRS voxel area identified a significant PCu/PCC cluster, confirming the positive correlation between scene-related BOLD activity and PCu/PCC tNAA. There were no correlations between PCu/PCC fMRI activity and Glx or GABA+ levels. These results demonstrate, for the first time, that scene activity in PCu/PCC is linked to local tNAA levels, identifying a neurochemical influence on inter-individual differences in the task-driven activity of a key brain hub.

## Introduction

The precuneus/posterior cingulate cortex (PCu/PCC) is a key brain hub and component of a core midline system, activated during autobiographical memory, prospection, contextual retrieval, spatial navigation, and mental scene construction (Bar et al., 2007; Buckner and Carroll, 2007; Hassabis and Maguire, 2007; van den Heuvel and Sporns, 2013; Andrews-Hanna et al., 2014). This system largely overlaps with the resting-state default-mode network (DMN), which is proposed to show greater activity during the resting state and deactivation during certain cognitive tasks [Fransson and Marrelec, 2008; Raichle et al., 2001; Raichle, 2015; Utevsky et al., 2014]. Based on its wider pattern of connectivity with medial temporal lobe regions, including parahippocampal and entorhinal cortices, plus the hippocampal formation [Hutchison et al., 2014; Kravitz et al., 2011; Parvizi et al., 2006; Passarelli et al., 2018], however, a recent proposal maintains that the PCu/PCC anchors a posteromedial or extended navigation system critical for the construction of “situation models” (comprising a particular time, place, and context) [Ranganath and Ritchey, 2012]. Such situation models support a diverse array of cognitive processes, underpinning episodic memory and future thinking, spatial navigation, and scene imagination, but also, importantly, online complex scene discrimination [Murray et al., 2017; Ranganath and Ritchey, 2012].

Much of our knowledge of the role of PCu/PCC in scene-related cognition is based on measurements tied to neuronal activity, including blood-oxygenation-level dependent (BOLD) functional magnetic resonance imaging (fMRI) in humans [Bzdok et al., 2015; Spreng et al., 2009] and single unit recording studies in both humans [Fox et al., 2018] and non-human primates [Dean, 2006; Sato et al., 2010]. Further evidence comes from neuropsychological investigations of scene-based processing following brain lesions [Futamura et al., 2018; Suzuki et al., 1998], including Alzheimer disease-linked neurodegeneration [Chan et al., 2016; Irish et al., 2015]. One avenue that remains substantially unexplored is the neuro-biochemical factors underpinning PCu/PCC activity during complex scene cognition. Given the importance of this core brain hub in healthy brain function as well as in disease (e.g. Alzheimer’s disease and psychiatric disorders such as schizophrenia [Leech and Sharp, 2014]), improved understanding of the biochemical mechanisms underpinning the PCu/PCC fMRI response could provide useful insight into the physiological basis of its activity, which could in turn provide insight into factors underpinning activity alterations in disease and disease risk [Duncan et al., 2014].

Neuro-biochemical relationships can be assessed by combining fMRI with proton magnetic resonance spectroscopy (^1^H-MRS), as this enables the investigation of the relationship between activity and local concentrations of metabolites non-invasively *in vivo* [Duncan et al., 2014]. The most commonly studied ^1^H-MRS metabolites are γ-amino-butyric acid (GABA) and glutamate (Glx), neurotransmitters which indicate inhibitory and excitatory tone, respectively, and N-acetyl-aspartate (tNAA), a neuronal marker associated with mitochondrial energy metabolism [Farrant and Nusser, 2005; Moffet et al., 2007; Rae, 2014; Stagg and Rothman, 2014]. A previous fMRI-MRS study on default-mode-related *deactivation* found that higher resting state PCu/PCC GABA+ and lower PCu/PCC Glx concentrations are associated with greater suppression of default-mode PCu/PCC activity during a working memory task [Hu et al., 2013]. In another study [Hao et al., 2013], higher tNAA was associated with reduced suppression of default-mode PCu/PCC activity during an auditory monitoring task. No study has yet assessed the relationship between task-driven *activation* in PCu/PCC and intrinsic regional levels of brain metabolites.

In this study, therefore, we investigated whether inter-individual differences in PCu/PCC activity elicited during complex scene discrimination would be related to inter-individual differences in resting state levels of PCu/PCC metabolites. An fMRI odd-one-out discrimination paradigm (“oddity”) [Hodgetts et al., 2015; Shine et al., 2015] was used to investigate PCu/PCC activity in response to different stimulus categories (scenes, faces, objects). First, we hypothesised there would be a greater PCu/PCC BOLD response during scene, compared to face or object, oddity, extending previous findings of a role for PCu/PCC in complex scene cognition to involvement in online perception [Ranganath and Ritchey, 2012; Spreng et al., 2009]. Second, ^1^H-MRS metabolites were measured in the PCu/PCC region plus in a control region in the occipital lobe (OCC), as in Hu et al., 2013. We hypothesised a positive relationship between PCu/PCC scene-related activity and tNAA levels (consistent with Hao et al., 2013), a positive relationship between PCu/PCC scene-related activity and Glx, and a negative relationship with GABA+ (consistent with Hu et al., 2013). Finally, we predicted there would be no relationship between PCu/PCC BOLD for complex scene discrimination and OCC metabolites; a finding that would be supportive of regional specificity of any PCu/PCC BOLD-MRS relationships [Duncan et al., 2014; Greenhouse et al., 2016; Hu et al., 2013].

## Methods

### Participants

Forty Cardiff University undergraduate and postgraduate students participated in this study (12 male, mean age 22.1 years, standard deviation 2.1 years, range 18.9-26.0 years). Participants were right-handed, had normal or corrected-to-normal vision, and no history of neurological or psychiatric disorders. The study received ethical approval from the Cardiff University School of Psychology Research Ethics Committee, and all participants provided written informed consent.

Due to the impact of menstrual cycle phase on ^1^H-MRS metabolite concentrations [Batra et al., 2008; Epperson et al., 2002], scans of female participants were scheduled during their luteal phase (days 15-28 of the cycle, where day 1 was defined as the first day of menstruation). As GABA+ concentration does not significantly differ between pill-on and pill-free days, no restrictions were placed on scan scheduling if female participants were taking the contraceptive pill [De Bondt et al., 2015].

### Overview of experimental design

Participants completed the perceptual odd-one-out fMRI task, an anatomical MRI scan, and four ^1^H-MRS scans within the same scan session.

### Perceptual odd-one-out (‘oddity’) fMRI task

The odd-one-out task was identical to that used in Shine et al. (2015) and Hodgetts et al. (2015). Participants were shown three stimuli from the same visual category on each trial and instructed to select the stimulus that was the ‘odd-one-out’ as quickly and as accurately as possible. Four stimuli categories were used: scenes (photographs of real-world outdoor scenes, e.g. parks, rivers, streets, buildings, taken by experimenters of Shine et al., 2015), faces (photographs of male and female faces, obtained from the Psychological Image Collection at Stirling (PICS) (http://pics.stir.ac.uk/), objects (real-world everyday items, e.g. televisions, chairs, kitchen implements, obtained from the Hemera Object Database, Vol. 1–3) or squares (taken from Barense, Henson, Lee, & Graham, 2010). For the scene, face and object categories, two of the images represented the same location, face or object, but were shown from a different viewpoint, and the third image was obtained from a visually similar, but unique, location, person or object. The squares acted as a baseline condition, with two of the squares being of equal size, while the third square differed in size, either slightly larger or smaller. Examples of a trial from each condition are shown in Figure 1A. Participants selected the odd-one-out by pressing the corresponding button on a 3-button MRI-compatible response box, held in their right hand. The position of the correct item was counterbalanced to appear in each location an equal number of times within each condition. All images were shown only once during the task (i.e. trial-unique).

**Figure 1:**
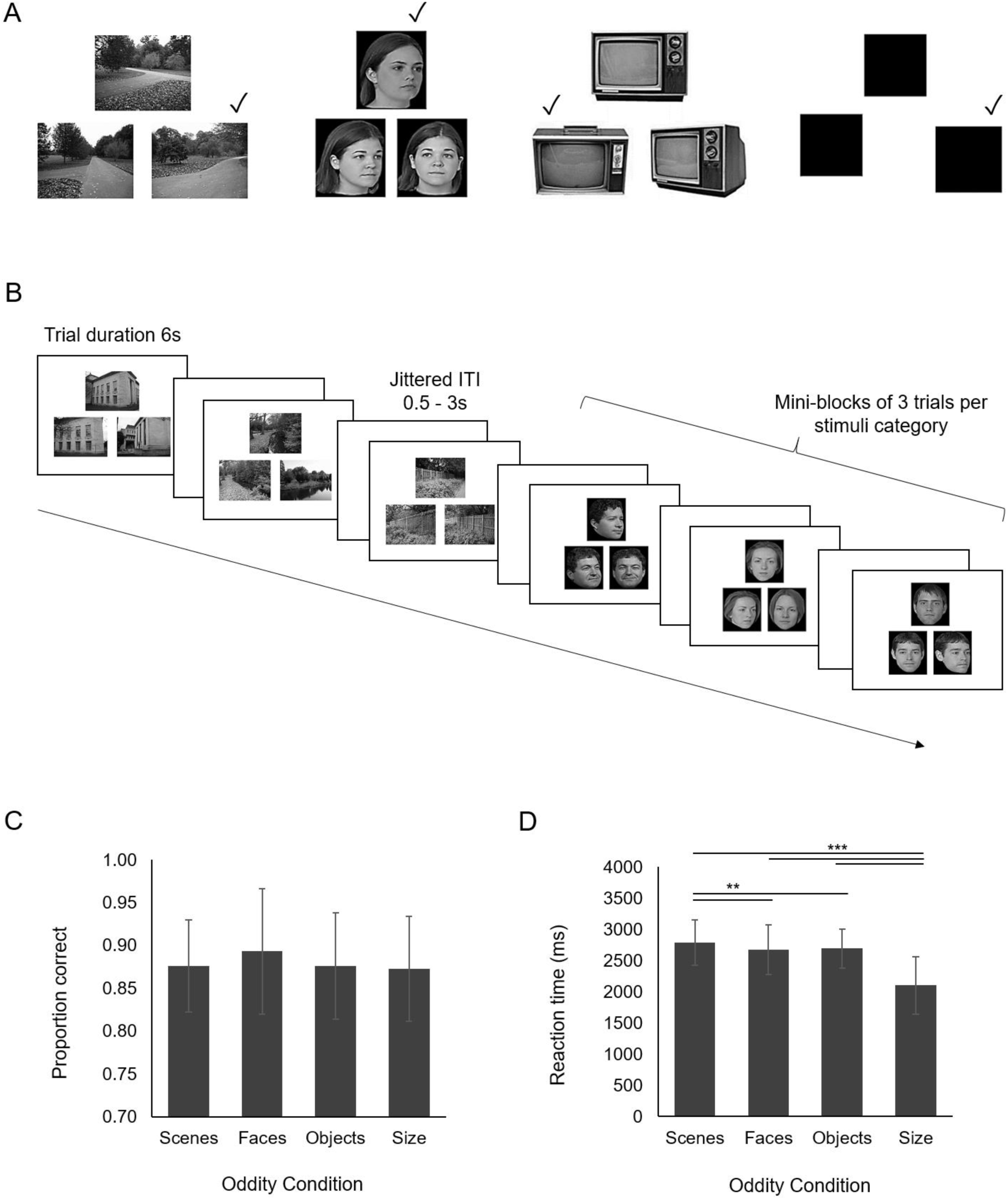
FMRI odd-one-out task. (A) Examples of a trial for each condition. Ticks indicate the correct odd-one-out. (B) Experimental design. Three trials from each category were presented in mini-blocks. Examples of a scene and face mini-block are shown. The order of mini-blocks was counterbalanced. (C and D) Behavioural results for the four stimulus categories (mean and standard error). N=35 (matched to the fMRI data shown below). (C) Proportion correct; (D) Reaction time. ^∗∗^ indicates statistically significant at p<0.01 and ^∗∗∗^ at p<0.001.

The experimental design is shown in Figure 1B. Each trial was presented for 6 seconds, and participants were required to make their response before the images disappeared from the screen. There was a jittered inter-trial interval of 500-3000ms, during which participants were presented with a blank white screen. Trials were arranged into mini-blocks of three trials per stimulus category, to reduce task-switching demands. The order of mini-block categories was counterbalanced between participants.

The experiment was divided into three fMRI runs, each consisting of 72 trials, and each run lasted 11 minutes. Eighteen trials per stimulus category per run were presented, resulting in a total of 54 trials per condition over the three runs. The experiment was implemented using E-prime version 2.0 (Psychology Software Tools, Inc., Sharpsburg, PA). The task was viewed in a mirror mounted on top of the head coil, which allowed participants to view a projector screen (Canon S×60 LCOS projector system combined with the Navitar SST300 zoom converter lens) located behind the scanner.

### MRI Acquisition

All scans were performed at the Cardiff University Brain Research Imaging Centre (CUBRIC) on a 3T General Electric (GE) HDx scanner fitted with an 8-channel phased array head coil. A 3D T1-weighted, fast spoiled gradient echo (FSPGR), structural MRI scan was obtained for each participant (TE/TR = 3.0/7.9ms; TI=450ms; flip angle 20° data matrix 256×192×176; field of view 256×192×176mm^3^; acquisition time approx. 7 minutes). The FSPGR was used to aid ^1^H-MRS voxel placement during scanning, and also as part of the subsequent fMRI and ^1^H-MRS data analysis.

### fMRI Acquisition

T2^∗^-weighted images were acquired using a gradient-echo, echo-planar imaging (EPI) sequence (TE/TR = 35/3000ms; flip angle 90°; field of view 220mm; data matri× 64×64). Each of the 3 fMRI runs consisted of 220 volumes, with each volume comprising 42 axial slices collected in a bottom-up interleaved order. The slice thickness was 2.8mm with an interslice gap of 1.0mm, which created an effective voxel size of 3.8×3.8×3.8mm^3^. Slices were aligned along the anterior commissure-posterior commissure (AC-PC) line, then a 30° axial-to-coronal tilt was introduced (in the ‘anterior up’ sense), to reduce signal dropout in the medial temporal lobe (MTL) caused by dephasing of the MRI signal due to nearby air-tissue and bone-tissue interfaces [Deichmann et al., 2003]. A field map was acquired to improve registration and reduce image distortion from magnetic-field inhomogeneity (TEs of 7ms and 9ms). Each fMRI run began with four dummy volumes, to allow the signal to reach T1 equilibrium prior to acquisition.

### ^1^H-MRS Acquisition

Single voxel proton spectra were acquired from the PCu/PCC (the voxel of interest, measuring 2×2×3cm^3^), and the occipital cortex (OCC, the control voxel, measuring 3×3×3cm^3^). Examples of voxel placement and landmarks used for placement are shown in Figure 2A. The landmarks were developed through a pilot study assessing test-retest reliability of voxel placement and metabolite concentrations. To place the PCu/PCC voxel, an odd number of AC-PC aligned slices were acquired from the bottom to the top of the corpus callosum (typically 5 or 7 slices), and the middle slice was chosen (shown by “mid” black line in Figure 2A). The PCu/PCC voxel was placed on top of this line corresponding to the middle slice, and adjusted to lie posterior to the ventricles to avoid placing the voxel in an area of CSF, which could introduce artefacts to the ^1^H-MRS spectrum. The occipital voxel was placed above the tentorium cerebelli (shown by arrow in Figure 2A) and adjusted so it did not contain any scalp tissue, which would have resulted in lipid contamination in the spectra.

**Figure 2:**
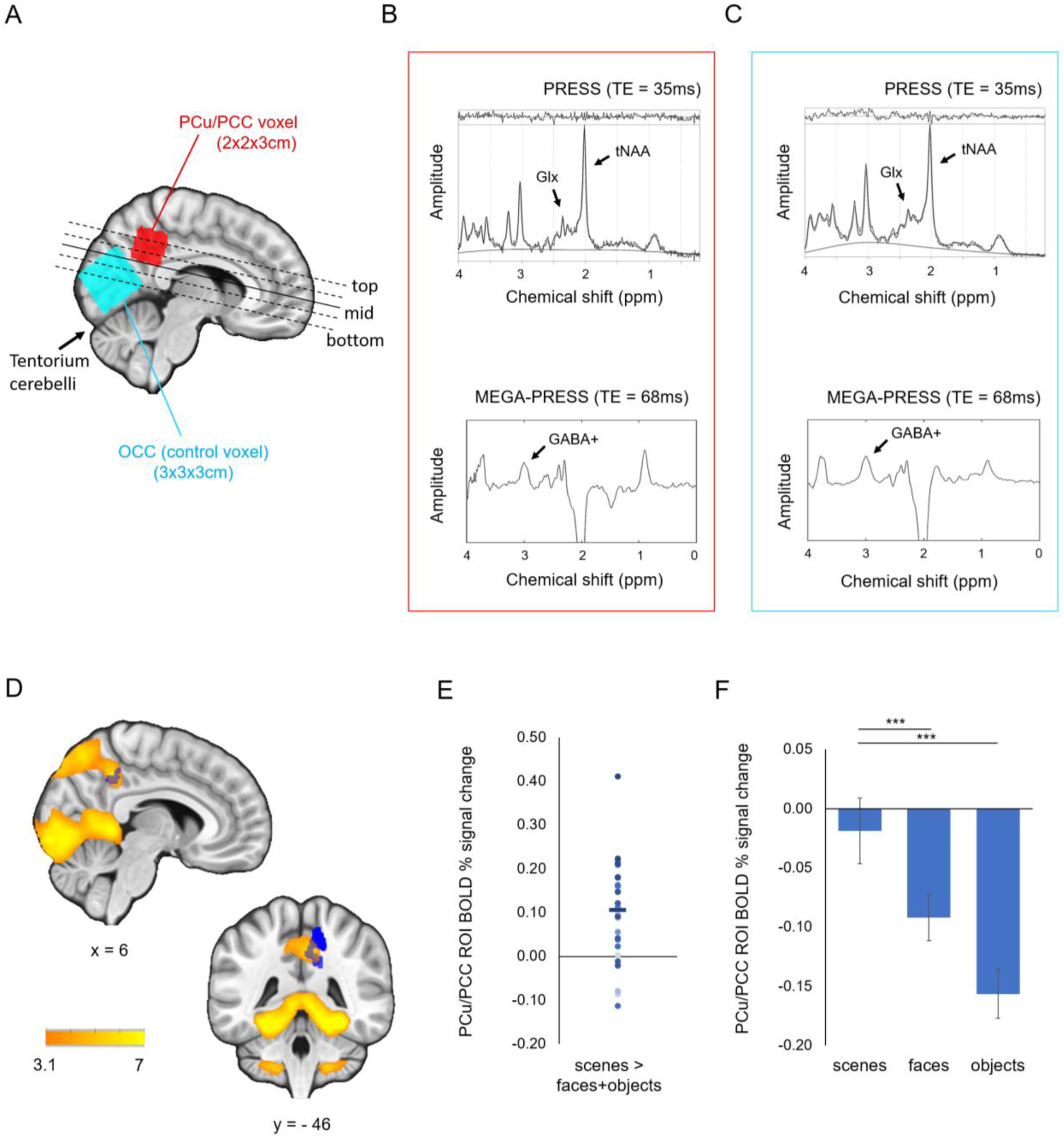
^1^H-MRS voxel placement and spectra and fMRI results. (A) ^1^H-MRS voxel placement for one participant. Voxels have been transformed into and overlaid on standard space. All brain data are rendered on the MNI152 2mm standard brain template. The dotted black lines represent slices acquired from the top to bottom of the corpus callosum, which were collected to identify the mid-slice (continuous black line) to use as a landmark for PCu/PCC voxel placement. The black arrow points to the tentorium cerebelli, used as the landmark line for OCC voxel placement. (B and C) Example of a PRESS and MEGA-PRESS spectrum from PCu/PCC voxel (B) and OCC voxel (C) in one participant. (D) FMRI results, showing significant whole brain clusters (n=35 participants) reflecting greater activity for scenes compared to faces or objects (Z>3.1, p<0.05). Blue shows the location of the *a priori* PCu/PCC ROI (from Shine et al., 2015). (E) BOLD percent signal change extracted from the *a priori* PCu/PCC ROI for the contrast scenes>faces+objects. Dots show each participant’s BOLD percent signal change for this contrast, and horizontal line shows the mean. (F) Mean PCu/PCC ROI BOLD percent signal change for each condition relative to baseline condition of size, ± standard error. ^∗∗∗^ indicates statistically significant at p<0.001.

In each voxel, one point-resolved spectroscopy (PRESS) scan was obtained to measure tNAA and Glx (TE/TR = 35/1500ms; number of averages = 128; scan time 4mins) [Bottomley, 1984]. GABA concentration (1-2mM) is very low in comparison to the other metabolites (2-15mM) and the peak resonances of GABA are of low amplitude due to J-coupling and are masked by the other metabolites (mainly by creatine at 3.0ppm) [Harris et al., 2017; Mullins et al., 2014]. Therefore, a spectral editing acquisition (Mescher-Garwood PRESS, MEGA-PRESS) [Mescher et al., 1998] was used to alleviate the difficulties in accurately quantifying GABA. MEGA-PRESS acquisitions include additional editing pulses placed symmetrically about the water resonance (4.7ppm) resulting in editing pulses at 1.9ppm (edit on) and at 7.5ppm (edit off) in order to subtract the creatine peak, enabling accurate GABA detection and quantification. One MEGA-PRESS acquisition per voxel was obtained to measure GABA + coedited macromolecules, “GABA+”, (TE/TR = 68/1800ms; OCC 166 edit on/off pairs, scan time 10mins; PCu/PCC 256 on/off pairs, scan time 15mins (longer scan used time in smaller PCu/PCC voxel to increase signal-to-noise)). Some studies have used the edit off scan to quantify tNAA and Glx, however here we used a separate PRESS scan because short TE PRESS scans (e.g. TE 35ms, rather than TE 68ms as in the MEGA-PRESS edit off scan) have been found to be more accurate and repeatable at quantifying these metabolites. For example, a test-retest reliability study of PCC glutamate found that over three intra-scan repeats, the 35ms PRESS scan produced a lower coefficient of variation and lower Cramer-Rao Lower Bounds (CRLBs) for Glx than PRESS scans with a longer TE [Hancu, 2009]. Shimming was performed before all ^1^H-MRS scans to ensure waterlinewidth of 10Hz or lower, in order to obtain sharp peaks in the resulting ^1^H-MRS spectrum.

### Data Analysis

#### Behavioural Data Analysis

Oddity task performance was assessed by calculating the accuracy (proportion of correct odd-one-out responses) and reaction time (RT, difference between stimulus onset and button press, in milliseconds, for correct trials only) on each of the four stimulus conditions. This was calculated for each participant and each experimental run separately. Values were averaged over the three runs to obtain a mean accuracy and RT measure for each condition.

#### fMRI Data Analysis

The fMRI data were pre-processed and analysed using FSL version 4.1, via the FMRI Expert Analysis Tool (FEAT) (http://www.fmrib.ox.ac.uk/fsl/). Pre-processing consisted of removal of the brain from the skull via the FSL brain extraction tool (BET) [Smith, 2002], motion correction using MCFLIRT [Jenkinson et al., 2002], spatial smoothing using a full-width-at-half-maximum (FWHM) Gaussian kernel of 5mm, mean-based intensity normalisation, high pass temporal filtering set to a cut-off value of 100 seconds, and field map unwarping of the EPI data using FUGUE [Jenkinson, 2003].

A general linear model (GLM), consisting of four explanatory variables (the four task conditions: scenes, faces, objects and size), was conducted for each subject and task run. An event-related design was used, and trials were modelled as 6s events from the start of the trial presentation. Trial start times, relative to scan onset, were obtained from each participant’s E-prime output file. As in previous studies, only correct trials were included in the GLMs [Hodgetts et al., 2017; Shine et al., 2015]. A parameter estimate image was created for each category relative to the size baseline condition (i.e. scenes>size, faces>size and objects>size), and for the planned contrast of scenes relative to the other main stimulus categories (scenes>faces+objects). Each run was examined for participant movement and excluded if movement exceeded one fMRI voxel (>3.8mm). This resulted in exclusion of two participants’ data, and one out of the three runs for five participants. The three experimental runs for each participant (and two experimental runs for five participants) were coregistered to the Montreal Neurological Institute (MNI152) 2mm template using FLIRT and combined using a fixed-effects model in FEAT.

To identify regions that showed a group-level preferential response to scenes compared to the other stimulus categories (scenes>faces+objects), a group-level analysis was performed using the FMRIB Local Analysis of Mixed Effects tool version 1 (FLAME 1; Beckmann et al., 2003; Woolrich et al., 2004). The resulting group-averaged statistical map was thresholded with a cluster-determining threshold of p = 0.001 [Eklund et al., 2016] with a familywise error-corrected cluster threshold of p = 0.05 based on Gaussian random fields theory.

To obtain a quantitative measure of BOLD percent signal change in the PCu/PCC (in order to correlate with ^1^H-MRS metabolite levels), an independently defined PCu/PCC region of interest (ROI) was used. The BOLD percent signal change in the PCu/PCC ROI for each contrast of scenes, faces and objects relative to size, and for scenes>faces+objects was extracted using the Featquery tool in FSL. The ROI chosen to sample PCu/PCC was a binarized mask of scene-related activity taken from Shine et al. (2015), which used the same fMRI paradigm, but in a different sample of individuals (peak voxel x=16, y= −46, z=30 in MNI co-ordinates, cluster size 273 voxels) (see Figure 2D).

#### ^1^H-MRS Data Analysis

PRESS data were analysed using TARQUIN (Totally Automatic Robust Quantification In NMR) version 4.3.3 [Wilson et al., 2011]. Since it is difficult to accurately separate the peak of NAA at 2.01ppm from that of N-acetyl-aspartyl-glutamate (NAAG) at 2.04ppm, these were combined to create a “total NAA” or tNAA measure (NAA+NAAG) (Rae, 2014). Similarly, it is difficult to accurately distinguish the spectra of glutamate and glutamine due to their largely overlapping resonances, so these measures were combined to create a composite glutamine + glutamate measure, or “Glx” [Rae, 2014; Stagg and Rothman, 2014]. To ensure good quality data, metabolites were excluded if the CRLB was above 20%, consistent with the exclusion criteria commonly found in the ^1^H-MRS literature [Cavassila et al., 2001; Hu et al., 2013; Stagg and Rothman, 2014]. This resulted in the exclusion of two PCu/PCC tNAA, three PCu/PCC Glx, one OCC tNAA and one OCC Glx measurements.

MEGA-PRESS data were analysed using GANNET (GABA-MRS Analysis Tool) version 2.0 [Edden et al., 2014]. The GABA concentration measured in the MEGA-PRESS scans represents GABA plus co-edited macromolecules, and is referred to as “GABA+” [Mullins et al., 2014]. Data quality was assessed by two raters (authors AGC and CJE) using a 3-point rating scale (as in Lipp et al., 2015). Ratings were done blind to the fMRI results. Based on this, eleven PCu/PCC GABA+ and one OCC GABA+ measurements were excluded, considered by the raters to be poor quality data-sets.

All metabolite concentrations were corrected for voxel composition: in each participant’s native space, PCu/PCC and OCC ^1^H-MRS voxels were segmented using the FAST tool in FSL to obtain a value for the proportion of cerebrospinal fluid (CSF), grey matter (GM) and white matter (WM) [Zhang et al., 2001]. All metabolites were quantified using H2O as an internal concentration reference and are expressed as a concentration in millimoles (mM) per unit tissue volume. The metabolite signals were corrected to account for the CSF fraction of the voxel (due to the negligible metabolite concentration within CSF) and the water reference signal was corrected to account for the differing water content of CSF, GM and WM (as in Harris et al., 2015a, 2015b; Lipp et al., 2015).

### Statistical Analysis

#### fMRI

A repeated-measures ANOVA, implemented in SPSS version 20 (SPSS Inc., Chicago, IL, US), was used to assess whether the PCu/PCC ROI percent signal change for the contrasts scenes>size, faces>size and objects>size were significantly different from each other. Greenhouse-Geisser correction was used where the assumption of sphericity was not met.

#### fMRI-MRS

Since there is no well-established benchmark method of testing the relationship between BOLD and ^1^H-MRS metabolites (for example, see Harris et al., (2015a) for five different analysis strategies), two complementary methods for assessing this relationship were implemented, consistent with Enzi et al. (2012) and Lipp et al. (2015):

##### (1) Pearson correlation of ^1^H-MRS metabolites and PCu/PCC ROI BOLD

Pearson correlations were used to assess the relationship between the BOLD response constrained to the independently defined fMRI ROI in the PCu/PCC for the contrast scenes>faces+objects and resting state tNAA, Glx and GABA+ concentrations quantified from the PCu/PCC ^1^H-MRS voxel. The independently defined ROI was chosen as it gives an unbiased estimate of effect size [Kriegeskorte et al., 2009], and is a smaller region than the large MRS voxel (fMRI ROI volume 2184mm^3^, PCu/PCC MRS voxel volume 12,000mm^3^) (as per Enzi et al., 2012). To assess the regional selectivity of any identified relationship (see Duncan, Wiebking, Munoz-Torres, & Northoff, 2013), Pearson correlations were performed between PCu/PCC ROI BOLD for scenes>faces+objects and OCC tNAA, Glx and GABA+. In total, this created two families of three correlations (one family of interest and one control family). Bonferroni correction was applied within each family (i.e. family-wise error correction: 0.05/3 = p 0.017). This provided correction for multiple comparisons, while avoiding an overly stringent Bonferroni correction, which would have had the potential for a high risk of false negative results (Nakagawa, 2004; also see http://daniellakens.blogspot.co.uk/2016/02/why-you-dont-need-to-adjust-you-alpha.html). All statistical tests were performed using SPSS version 20 (SPSS Inc., Chicago, IL, US).

To test whether correlations between PCu/PCC BOLD and metabolite concentrations in the PCu/PCC and OCC voxels were statistically different from each other, a directional Olkin’s Z test, testing if the correlation with PCu/PCC tNAA was stronger than OCC tNAA, was performed as implemented in the “cocor” R package (Diedenhofen and Musch, 2015; http://comparingcorrelations.org).

As recommended by Dienes and McLatchie (2017), Default Bayes factors (and 95% Bayesian credibility intervals) were also computed for all BOLD-MRS correlations, using JASP version 0.8.1.12 (https://jasp-stats.org/). The Bayes factor (BF) grades, on a continuous scale, the strength of the evidence that the data provide either for the alternative hypothesis (H1) versus the null (H0) expressed as BF_10_, or for the null hypothesis versus the alternative hypothesis expressed as BF_01_ [Jarosz and Wiley, 2014; Wetzels and Wagenmakers, 2012]. A BF_10_ or BF_01_ of 1 indicates that the observed finding is equally likely under H0 and H1. A BF_10_ much greater than 1 allows us to conclude that there is substantial evidence for H1 over H0. BF_10_ values substantially less than 1 provide strong evidence in favour of H0 over H1. Conversely a BF_01_ of much greater than 1 indicates there is strong evidence in favour of H0 over H1, whereas a BF_01_ of much less than 1 indicates strong support for H1 rather than H0. Where the Pearson correlations were significant, directional BFs for these correlations were expressed as BF_+0_ in favour of a positive correlation, or BF_-0_ for a negative correlation [Marsman and Wagenmakers, 2017]. Where the Pearson correlations were non-significant, BF_01_ in favour of the null (zero) correlation were computed.

##### (2) Voxel-wise GLM using ^1^H-MRS as a regressor

Three GLMs were created for the contrast of scenes>faces+objects, where each model included demeaned values of either PCu/PCC tNAA, Glx or GABA+ concentrations as a regressor. These analyses were restricted to a group PCu/PCC ^1^H-MRS voxel mask area (as in Enzi et al., 2012). Contrasts were set up so that results would show any voxels within this group mask that were positively or negatively correlated with concentrations of each metabolite. The group PCu/PCC ^1^H-MRS voxel mask was created in MNI space by combining all participants’ PCu/PCC MRS voxel masks, such that any MNI voxel present in any participant’s MRS voxel mask would be included in the group voxel mask (see Figure 5 for tNAA mask). The number of participants used to create the group masks differed for the tNAA, Glx and GABA+ analyses, as there were different numbers of participants remaining for these analyses after exclusions following data quality control (tNAA n=33, Glx n=32, GABA+ n=24).

The statistical threshold for the fMRI analysis was voxel-wise, uncorrected at p<0.01 (as in Hodgetts et al., 2015). To prevent false positives due to multiple-comparisons (as there were approximately 4000 fMRI voxels within each PCu/PCC MRS metabolite group mask), Monte-Carlo simulation was used to determine whether the size of any resulting cluster was statistically significant, using the 3dClustSim command in AFNI (2016 version, in which the bug identified in Eklund et al. (2016) had been fixed, https://afni.nimh.nih.gov/pub/dist/doc/program_help/3dClustSim.html). A cluster-corrected threshold of p<0.01 was selected (as in Hodgetts et al., 2015), which determined that any cluster larger than 42 voxels was statistically significant at p<0.01 for the tNAA mask and Glx mask, and 40 voxels for the GABA+ mask.

## Results

### Behaviour

There were no significant differences in the proportion of correct responses between the four conditions of the oddity task (F(3,102)= 1.16, p= 0.33, **η**^2^ _*p*_= 0.09), suggesting that task difficulty was matched across conditions (see Figure 1C). There was, however, a significant effect of condition on reaction time (F(3,102)= 97.25, p= 8.46×10^−30^, **η**^2^ _*p*_= 0.86). Post-hoc tests revealed reaction times were faster for the baseline condition of size than for scenes, face and objects (respectively: t(34)= 14.34, p= 5.64×10^−16^, Cohen’s d_av_= 1.65; t(34)= 11.13, p= 7.07×10^−13^, Cohen’s d_av_= 1.32 and t(34)= 11.71, p= 1.75×10^−13^, Cohen’s d_av_= 1.50). Reaction times were also slightly slower for scenes than for faces or objects respectively (scenes: t(34)= 3.02, p= 0.005, Cohen’s d_av_= 0.30; faces: t(34)= 2.83, p= 0.008, Cohen’s d_av_= 0.29) (see Figure 1C).

### fMRI

#### Whole brain results: PCu/PCC shows a greater BOLD response for scenes than faces and objects

The final fMRI sample size was 35, following exclusions due to scanner errors (n=3) and movement (n=2 + 1/3 of runs excluded for 5 participants). Figure 2D shows the whole brain activation map for the contrast of scenes compared to faces and objects. There was a significantly greater BOLD response in the PCu/PCC region for scenes>faces+objects, suggesting this region has relative scene selectivity (peak activation in MNI co-ordinates x= 6, y= −46, z= 42). Significant clusters were also identified for this contrast bilaterally in the parahippocampal gyrus, lateral occipital sulcus, lingual gyrus, inferior regions of the precuneus, temporal occipital fusiform cortex, and occipital pole (see Table 1), consistent with previous studies of scene-related activity [Hodgetts et al., 2016].

**Table 1:** Whole brain fMRI results. Clusters that showed significantly greater activity for the contrast of scenes>faces+objects. Brain region labels were defined using the Harvard-Cortical atlas tool in FSL.

| Brain region | Hemisphere | Z value | MNI co-ordinates of peak voxel |  |  |
| --- | --- | --- | --- | --- | --- |
|  |  |  | x | y | z |
| Temporal occipital fusiform cortex/ Lingual gyrus | Right | 8.81 | 30 | -48 | -12 |
| Temporal occipital fusiform cortex/ Lingual gyrus | Right | 8.69 | 30 | -52 | -12 |
| Temporal occipital fusiform cortex/ Lingual gyrus | Left | 8.59 | -28 | -52 | -10 |
| Precuneus/ Intracalcarine cortex/ Lingual gyrus | Left | 8.58 | -14 | -60 | 6 |
| Lingual gyrus / Precuneus | Right | 8.47 | 18 | -54 | 4 |
| Precuneus/Posterior cingulate cortex | Right | 6.34 | 6 | -46 | 42 |
| Posterior temporal fusiform cortex/ Temporal occipital fusiform cortex | Left | 8.41 | -26 | -42 | -16 |
| Superior/middle frontal gyri | Right | 4.71 | 28 | 2 | 60 |
|  | Left | 4.47 | -28 | 8 | 54 |

#### PCu/PCC ROI results: Greater BOLD percent signal change in ROI for scenes than faces or objects

The *a priori* PCu/PCC ROI overlapped with the PCu/PCC cluster identified in the whole brain analysis (see Figure 2D). Consistent with the whole brain findings, within the PCu/PCC ROI there was significantly greater activity for scenes>faces+objects (see Figure 2E for inter-individual variability in values). In support of this, when each condition was contrasted to the baseline condition of size, a repeated-measures ANOVA determined there was a significant effect of oddity condition on the BOLD percent signal change measured in the PCu/PCC ROI (F(1.42, 48.21)= 34.68, p= 1.6×10^−8^, **η**^2^ _*p*_= 0.51 (Greenhouse-Geisser corrected degrees of freedom)). Planned comparisons confirmed that this reflected significantly higher PCu/PCC activity for scenes compared to the face and object conditions when contrasted with size: scenes>size versus faces>size (F(1, 34)= 14.92, p= 4.8×10^−4^, **η**^2^ _*p*_= 0.31), and scenes>size versus objects>size (F(1, 34)= 52.65, p= 2.1×10^−8^, **η**^2^ _*p*_= 0.61) (see Figure 2F).

### ^1^H-MRS

Figure 2A shows a representative PCu/PCC and OCC voxel placement. Figure 2B and 2C show examples of each type of spectra obtained in each voxel from a single subject. Table 2 gives the mean metabolite concentrations in each voxel, and sample sizes remaining after data exclusions due to ^1^H-MRS and fMRI data quality assessment, to show data remaining for fMRI-MRS correlations.

**Table 2:** ^1^H-MRS results. Sample sizes given are those after data has been removed due to scanner errors, fMRI data quality assessment, and ^1^H-MRS quality control measures. Metabolites are expressed as a concentration in millimoles (mM) per unit tissue volume.

|  | PCu/PCC voxel |  |  | OCC voxel |  |  |
| --- | --- | --- | --- | --- | --- | --- |
|  | tNAA | Glx | GABA+ | tNAA | Glx | GABA+ |
| Sample size (from original n=40) | 33 | 32 | 24 | 34 | 34 | 34 |
| Mean concentration (mM) | 16.39 | 20.93 | 1.85 | 13.42 | 21.64 | 1.97 |
| Standard deviation | 0.90 | 2.87 | 0.27 | 1.40 | 2.75 | 0.19 |

### MRS-fMRI relationships

#### (1) Pearson correlation of PCu/PCC ROI BOLD with ^1^H-MRS metabolites

There was a significant positive correlation between the BOLD percent signal change for scenes>faces+objects in our *a priori* PCu/PCC ROI and the concentration of tNAA in the PCu/PCC ^1^H-MRS voxel (r(33)= 0.41, p= 0.017, BF_+0_ = 6.62, 95% Bayes CI = 0.09 and 0.65) (see Figure 3A).

**Figure 3:**
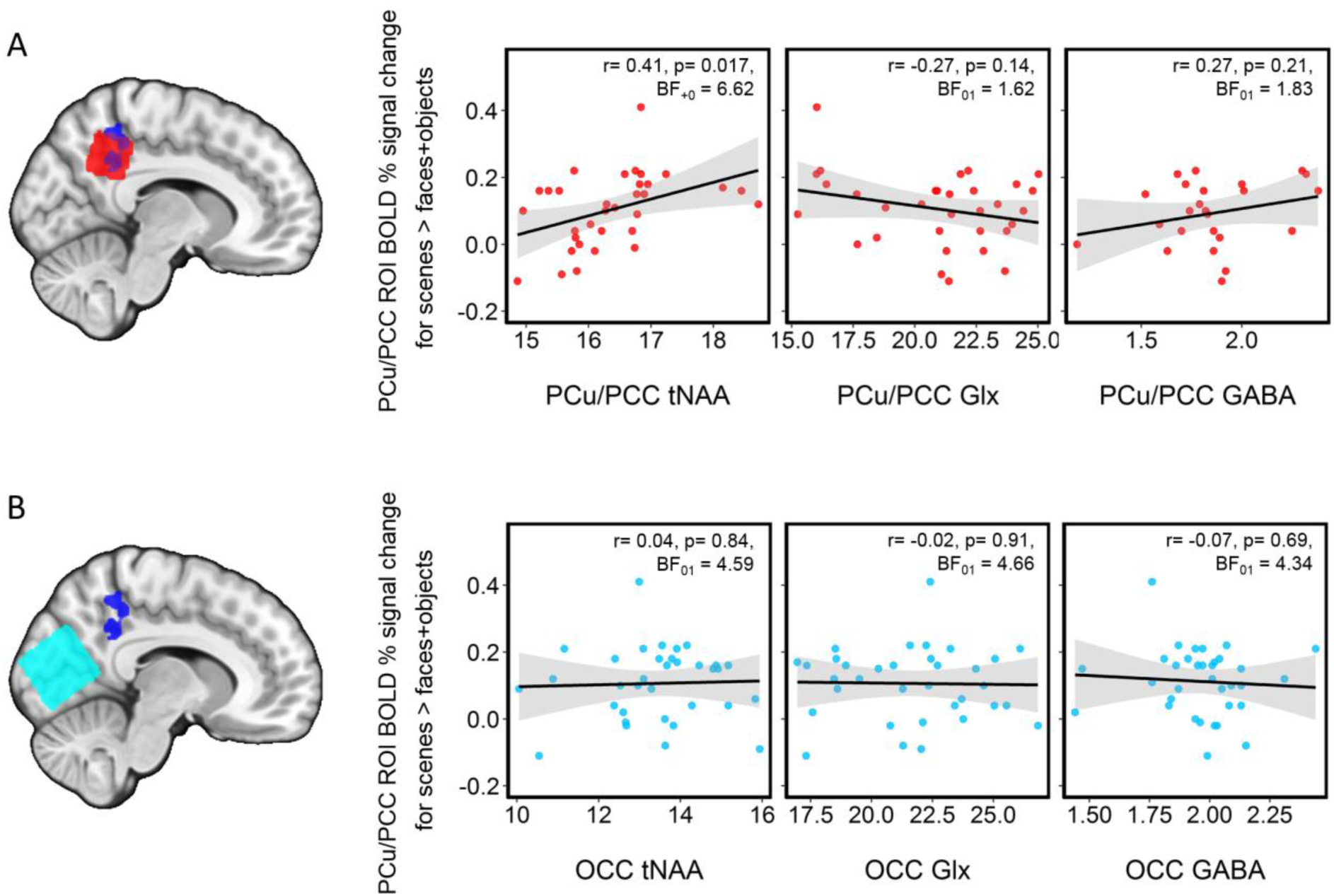
Pearson correlations of PCu/PCC ROI BOLD percent signal change for the contrast scenes>faces+objects with each of the three ^1^H-MRS metabolites from (A) the PCu/PCC MRS voxel (red data points) and (B) the OCC control voxel (blue data points). Metabolites are expressed as a concentration in millimoles (mM) per unit tissue volume. Grey shaded areas show the 95% confidence intervals.

This relationship appeared to be regionally selective, as there was no significant correlation between PCu/PCC BOLD for scenes>faces+objects and OCC tNAA (r(34)= 0.04, p= 0.84, BF_01_ = 4.59, 95% Bayes CI = −0.29 and 0.36) (see Figure 3B). The difference between correlations for PCu/PCC BOLD for scenes>faces+objects with PCu/PCC tNAA versus with OCC tNAA showed a trend towards being significant (Z = 1.58, p = 0.058).

There were no significant correlations between PCu/PCC BOLD for scenes>faces+objects and either PCu/PCC Glx (r(32)= −0.27, p=0.14, BF_01_ = 1.62, 95% Bayes CI = −0.54 and 0.09), or PCu/PCC GABA+ (r(24)= 0.27, p= 0.21, BF_01_ = 1.83, 95% Bayes CI = −0.14 and 0.58) (see Figure 3A). There was also no relationship between PCu/PCC BOLD for scenes>faces+objects with OCC Glx (r(34)= −0.02, p= 0.91, BF_01_ = 4.66, 95% Bayes CI = −0.34 and 0.31) or with OCC GABA+ (r(34)= −0.07, p= 0.69, BF_01_ = 4.34, 95% Bayes CI = −0.38 and 0.26) (see Figure 3B).

To confirm that the positive correlation identified between the PCu/PCC BOLD response for scenes and PCu/PCC tNAA was indeed driven by *activation* for scenes, rather than *deactivation* for faces and objects relative to scenes (as depicted in Figure 2F), additional correlations were performed between PCu/PCC BOLD for scenes>size baseline and PCu/PCC tNAA or OCC tNAA. Consistent with the above correlations, a significant positive relationship was identified between PCu/PCC BOLD for scenes>size and PCu/PCC tNAA (r(33)= 0.42, p= 0.016, BF_10_ = 7.03, 95% Bayes CI = 0.09 and 0.65). Again, this was regionally selective, as there was no significant correlation between PCu/PCC BOLD for scenes>size with OCC tNAA (r(34)= 0.09, p= 0.63, BF_01_ = 4.18, 95% Bayes CI = −0.25 and 0.40) (see Figure 4B). The difference between these two correlations showed a trend towards being significant (Z = 1.41, p = 0.079).

**Figure 4:**
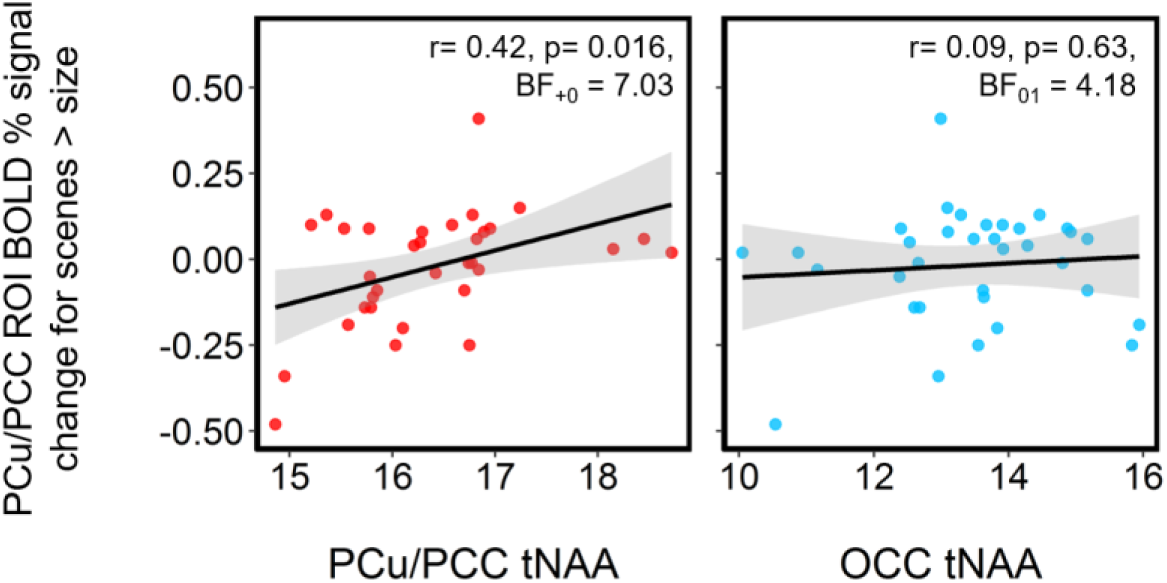
Pearson correlations of PCu/PCC ROI BOLD percent signal change for the contrast scenes>size with PCu/PCC or OCC tNAA. Metabolites are expressed as a concentration in millimoles (mM) per unit tissue volume. Grey shaded areas show the 95% confidence intervals.

#### (2) Voxel-wise GLM using ^1^H-MRS as a regressor

The relationship between PCu/PCC tNAA and BOLD for scenes>faces+objects identified above was also identified when PCu/PCC tNAA was used as a regressor in the GLM within the PCu/PCC ^1^H-MRS voxel mask area (see Figure 5). This revealed a significant cluster of 167 voxels where PCu/PCC tNAA was positively correlated with the BOLD response to scenes>faces+objects (peak MNI co-ordinate 8, −48, 48; peak Z=3.76). As depicted in Figure 5, this was located in a highly similar region to the *a priori* PCu/PCC ROI, supporting the selective relationship between PCu/PCC BOLD for scenes and PCu/PCC tNAA concentration.

**Figure 5:**
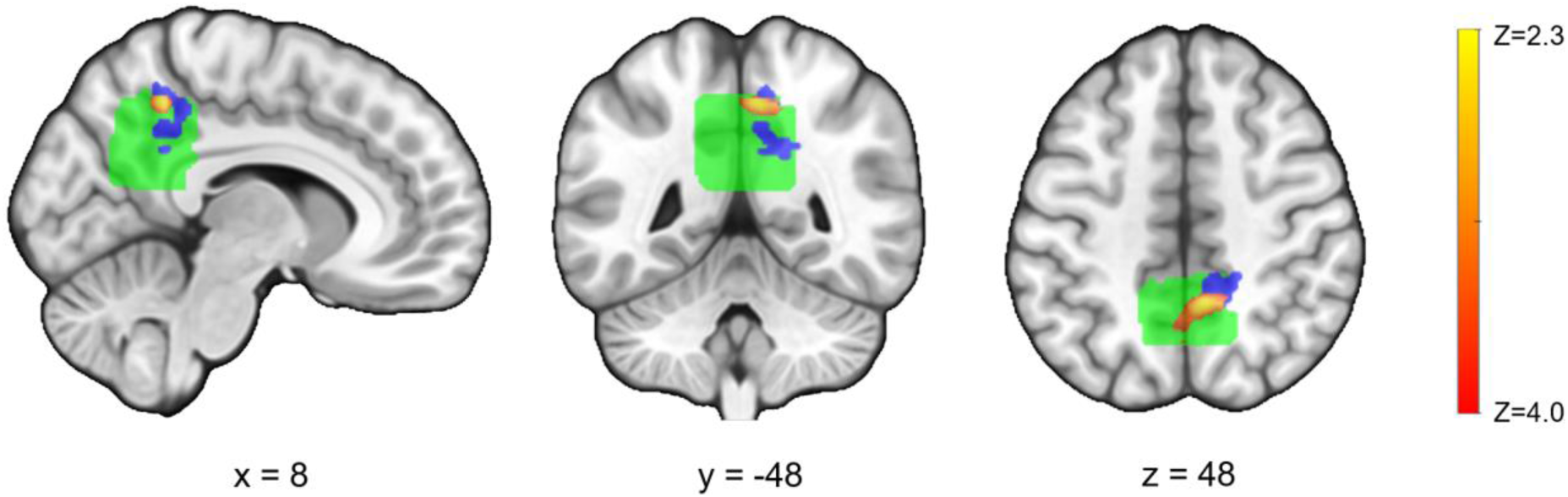
Voxel-wise regression analysis rendered on MNI152 standard brain template. Green shows the PCu/PCC ^1^H-MRS voxel mask area for the tNAA analysis, created by overlaying the participants’ PCu/PCC ^1^H-MRS voxels. Blue shows the *a priori* PCu/PCC mask used in the ROI analysis. Red/yellow shows the significant cluster of 167 voxels within the green ^1^H-MRS voxel mask area where PCu/PCC tNAA is positively correlated with BOLD for scenes>faces+objects (voxel-wise threshold of Z<2.3, cluster-wise threshold of p<0.01 corresponding to clusters > 42 voxels).

When Glx or GABA+ were included as regressors in separate GLMs, no significant clusters were identified reflecting positive or negative associations with PCu/PCC BOLD for scenes>faces+objects.

## Discussion

We adopted a multimodal functional and biochemical imaging approach to test whether individual differences in PCu/PCC activity during complex scene discrimination were linked to concentrations of PCu/PCC ^1^H-MRS metabolites. We found a significant positive correlation between the PCu/PCC BOLD response to scenes and the resting state concentration of tNAA in PCu/PCC. This pattern was not evident with Glx or GABA+ and was regionally specific to PCu/PCC, since OCC metabolite concentrations were not linked with PCu/PCC BOLD activity.

The PCu/PCC is a core posterior hub of the DMN [Fransson and Marrelec, 2008; Raichle, 2015; Utevsky et al., 2014]. Historically, the DMN was identified as a task-negative network, since it was reliably engaged during passive conditions, including fixation (Raichle et al., 2001). Far from being passive however, default activity during fixation is hypothesized to reflect spontaneous cognitive processes including autobiographical recall, imagination and prospection, although accounts differ as to the role of PCu/PCC in such processes [Andrews-Hanna et al., 2014; Buckner and Carroll, 2007; Hassabis and Maguire, 2007; Schacter et al., 2012; Spreng, 2012]. Here, both the ROI and whole brain analyses revealed that PCu/PCC responded significantly more to scenes than to either faces or objects, a finding inconsistent with a simple task-negative account of PCu/PCC function.

Our finding that dorsal PCu/PCC is particularly involved in complex scene discrimination is, however, consistent with other studies showing a role for activation in this region in a variety of scene-related tasks including scene working memory [Hodgetts et al., 2016]; mental scene construction [Hassabis et al., 2007; Palombo et al., 2018]; as well as large-scale spatial navigation in virtual reality environments [Ekstrom et al., 2017]. The PCu/PCC region is heavily interconnected with core scene processing regions including retrosplenial and parahippocampal cortices [Hutchison et al., 2014; Kravitz et al., 2011; Parvizi et al., 2006; Passarelli et al., 2018] (which were preferentially activated here by scenes alongside PCu/PCC), and also links the DMN with the dorsal attention network [Spreng et al., 2013], placing it in a key anatomical position to coordinate processes important for scene discrimination performance.

Our fMRI findings thus lend support to accounts in which PCu/PCC comprises a hub within a posteromedial system critical for the construction of mental scenes [Hassabis and Maguire, 2007] or more generally situation models [Murray et al., 2017; Ranganath and Ritchey, 2012], which critically underpin spatial navigation, autobiographical memory retrieval, simulation of future episodes and, as shown here, view-invariant scene perception. Successful performance on scene oddity depends on such internal models, because participants must be able to locate each viewpoint (and therefore reject the odd-one-out) within an overarching unified context, whose unseen aspects must be internally generated.

### BOLD-tNAA relationship

A key novel finding from this study was the positive association, across two complementary analysis approaches, between scene-selective PCu/PCC activity and intra-regional tNAA concentration. Specifically, where an individual showed a greater PCu/PCC BOLD response for scenes, this was associated with higher tNAA levels. Based on the proposed functions of tNAA, we offer 3 potential explanations for our findings:

i. NAA, which is present predominantly in neuronal cells [Simmons et al., 1991], is considered as a biomarker of neuronal and axonal density and integrity [Duarte et al., 2012; Rae, 2014]. The positive correlation between tNAA and the local BOLD response, therefore, could indicate that a higher density of neurons that respond to scenes (vs. faces and objects) leads to an increased scene BOLD response.
ii. NAA is synthesised in neuronal mitochondria (the site of aerobic respiration to produce ATP) [Patel and Clark, 1979], and there is evidence that NAA and mitochondrial function are coupled, as pharmacological inhibition of mitochondrial respiratory chain enzymes required for ATP synthesis are associated with a decrease in NAA [Bates et al., 1996]. ATP production is required following neuronal activity to replenish energy stores expended during this energy-demanding process [Logothetis, 2002]. The BOLD response is an indirect measure of the increase in nutrients (e.g. oxygen and glucose) delivered to the site of neuronal activity for ATP production. A second possible mechanism linking tNAA with BOLD, therefore, is that a higher BOLD response could represent greater functional capacity of the neuronal mitochondria to produce ATP, which is associated with levels of NAA.
iii. A third possible mechanism is based on NAA being a precursor molecule for N-acetyl-aspartyl-glutamate (NAAG) [Moffet et al., 2007; Rae, 2014]. In most ^1^H-MRS studies, NAA and NAAG are combined to create a composite tNAA measure [Rae, 2014]. A recent functional ^1^H-MRS (fMRS) study, however, assessed NAA and NAAG separately, and found that NAAG increases and NAA decreases in concentration following neuronal stimulation (Landim et al., 2016). NAAG induces a vasodilatory response in local blood vessels, via being exported from neurons to astrocytes where it binds to the mGluR3 glutamate receptor. This results in the release of vasoactive agents, which is in turn associated with an increase in the BOLD response [Baslow et al., 2016]. Further evidence that NAAG is associated with BOLD comes from the pharmacological inhibition of NAAG breakdown, which causes an increase in the BOLD response for a few minutes [Baslow et al., 2005]. Returning to the tNAA-BOLD relationship identified here, if there was a higher amount of regional tNAA (i.e. combined NAA and NAAG), there could be more substrate (NAA) to produce more NAAG. In turn, this could result in a greater vasoactive response, reflected in a greater BOLD response. Future (invasive) studies will be required to untangle these possible mechanisms. The causes of inter-individual variation in tNAA in young adulthood are not fully understood, but likely include a complex interplay between genetic and environmental factors over the lifecourse. Levels of tNAA have been shown to be highly heritable [Batouli et al., 2012], but are also sensitive to early environmental influences, including stress [McLean et al., 2012]. Future large scale studies of genetic and environmental influences on PCu/PCC metabolite levels in humans across the lifespan will be required to address this critical question, which may have implications for understanding lifespan influences on cognitive and neural aging [Jagust and Mormino, 2011].

### BOLD-GABA+ and BOLD-Glx relationships

We found no significant correlation between PCu/PCC BOLD scene activity and regional GABA+ or Glx, which contrasts with a previous study of DMN deactivation [Hu et al., 2013]. The lack of BOLD-Glx and BOLD-GABA+ correlations could reflect a lower signal-to-noise ratio for the multiple peaks of Glx and GABA+ which overlap with other metabolites, compared to the large peak of tNAA [Harris et al., 2017; Rae, 2014; Stagg and Rothman, 2014]. The data quality for tNAA was also better than that of GABA+, as many more GABA+ than tNAA measurements were excluded through quality control assessments. An alternative, more mechanistic reason could be that BOLD-tNAA and BOLD-GABA+ or BOLD-Glx relationships are capturing quite different physiological processes. For example, GABA+ and Glx could indicate inhibitory and excitatory tone [Farrant and Nusser, 2005; Rae, 2014; Stagg et al., 2011], whereas tNAA could reflect neuro-vascular coupling, as speculated above. In the absence of strong Bayes Factors in support of the null for BOLD-GABA+ and BOLD-Glx relationships, however, this possibility must remain speculative.

### Limitations

Although ^1^H-MRS has the advantage that it is the only neuroimaging technique that can contribute knowledge of biochemistry non-invasively *in vivo*, there are limitations to this technique. The large voxel sizes required to quantify metabolites present at very low concentrations limits the spatial resolution that ^1^H-MRS can provide in comparison to the high spatial resolution of fMRI [Stagg and Rothman, 2014]. This is particularly problematic for GABA quantification [Mullins et al., 2014]. An attempt was made at reducing the size of the PCu/PCC ^1^H-MRS voxel from the standard 3×3×3cm to 2×2×3cm to improve regional specificity in this study; this came, however, at the cost of several PCu/PCC MEGA-PRESS spectra being of too poor quality to use the GABA+ measurement obtained from this region. Moving to stronger magnetic field strengths (e.g. 7T) could improve signal-to-noise for the metabolite measurements, making it possible to reduce the voxel sizes to improve the spatial resolution of the ^1^H-MRS measures [Terpstra et al., 2016; Tkáč et al., 2009].

As is common in previous fMRI-MRS studies, a limitation of our approach was that the ^1^H-MRS and fMRI scans were acquired separately. The concentrations of tNAA, Glx and GABA+ measured, therefore, reflected their concentrations during an unconstrained resting state rather than during the task. Recent fMRI-MRS studies are now attempting to assess task-related ^1^H-MRS changes, through functional MRS (fMRS) [Apšvalka et al., 2015; Landim et al., 2016], or through collecting ^1^H-MRS and fMRI data simultaneously, by acquiring both within the same TE [Ip et al., 2017]. Future studies could assess the relationship between task-related tNAA (and other metabolites) and the BOLD response in PCu/PCC.

### Summary

In conclusion, our multimodal imaging study revealed that dorsal PCu/PCC shows a selective response to scenes compared to faces or objects. This is consistent with the view that this region is a key hub of a posteromedial network critical for scene or situation model construction, including in the service of online perception. The magnitude of the PCu/PCC BOLD response to scenes (relative to the other stimulus categories, including low-level baseline) was positively associated with the concentration of the PCu/PCC ^1^H-MRS metabolite tNAA, and not with concentrations of PCu/PCC GABA+ or Glx, nor with OCC metabolite concentrations. This finding provides novel evidence on the biochemical correlates of PCu/PCC activity, raising the intriguing possibility that neuronal density and/or mitochondrial energy metabolism could underpin individual differences in the magnitude of the scene-selective BOLD response in this key brain hub.

## Acknowledgements

We are grateful to Mark Mikkelsen for sharing Matlab scripts for ^1^H-MRS analysis, and to Peter Hobden and Martin Stuart for assistance with scanning. Thank you also to Jon Shine and Morgan Barense for stimuli creation. The work was funded by the Wellcome Trust (Strategic Award – KSG, ADL and CJH, 104943/Z/14/Z) and Medical Research Council (KSG and CJH, G1002149). AGC’s PhD studentship was funded by the Cardiff University Neuroscience and Mental Health Research Institute.

## References

Andrews-Hanna JR, Saxe R, Yarkoni T (2014): Contributions of episodic retrieval and mentalizing to autobiographical thought: Evidence from functional neuroimaging, resting-state connectivity, and fMRI meta-analyses. Neuroimage 91:324–335. https://doi.org/10.1016/j.neuroimage.2014.01.032.

Apšvalka D, Gadie A, Clemence M, Mullins PG (2015): Event-related dynamics of glutamate and BOLD effects measured using functional magnetic resonance spectroscopy (fMRS) at 3 T in a repetition suppression paradigm. Neuroimage 118:292–300. https://doi.org/10.1016/j.neuroimage.2015.06.015.

Bar M, Aminoff E, Mason M, Fenske M (2007): The units of thought. Hippocampus 17:420–428. http://doi.wiley.com/10.1002/hipo.20287.

Barense MD, Henson RNA, Lee ACH, Graham KS (2010): Medial temporal lobe activity during complex discrimination of faces, objects, and scenes: Effects of viewpoint. Hippocampus 20:389–401.

Baslow MH, Cain CK, Sears R, Wilson DA, Bachman A, Gerum S, Guilfoyle DN (2016): Stimulation-induced transient changes in neuronal activity, blood flow and N-acetylaspartate content in rat prefrontal cortex: a chemogenetic fMRS-BOLD study. NMR Biomed 29:1678–1687. https://doi.org/10.1002/nbm.3629.

Baslow MH, Dyakin V V, Nowak KL, Hungund BL, Guilfoyle DN (2005): 2-PMPA, a NAAG Peptidase Inhibitor, Attenuates Magnetic Resonance BOLD Signals in Brain of Anesthetized Mice: Evidence of a Link Between Neuron NAAG Release and Hyperemia. J Mol Neurosci 26:001–016. http://link.springer.eom/10.1385/JMN:26:1:001.

Bates TE, Strangward M, Keelan J, Davey GP, Munro PM, Clark JB (1996): Inhibition of N-acetylaspartate production: implications for 1H-MRS studies in vivo. Neuroreport 7:1397–1400.

Batouli SAH, Sachdev PS, Wen W, Wright MJ, Suo C, Ames D, Trollor JN (2012): The heritability of brain metabolites on proton magnetic resonance spectroscopy in older individuals. Neuroimage 62:281–289. https://doi.org/10.1016/j.neuroimage.2012.04.043.

Batra NA, Seres-Mailo J, Hanstock C, Seres P, Khudabux J, Bellavance F, Baker G, Allen P, Tibbo P, Hui E, Le Melledo J-M (2008): Proton Magnetic Resonance Spectroscopy Measurement of Brain Glutamate Levels in Premenstrual Dysphoric Disorder. Biol Psychiatry 63:1178–1184. https://doi.org/10.1016/j.biopsych.2007.10.007.

Beckmann CF, Jenkinson M, Smith SM (2003): General multilevel linear modeling for group analysis in FMRI. Neuroimage 20:1052–1063. https://doi.org/10.1016/S1053-8119(03)00435-X.

De Bondt T, De Belder F, Vanhevel F, Jacquemyn Y, Parizel PM (2015): Prefrontal GABA concentration changes in women—Influence of menstrual cycle phase, hormonal contraceptive use, and correlation with premenstrual symptoms. Brain Res 1597:129–138. http://dx.doi.org/10.1016/j.brainres.2014.11.051.

Bottomley (1984): Selective volume method for performing localized NMR spectroscopy. US Patent 4480228.

Buckner RL, Carroll DC (2007): Self-projection and the brain. Trends Cogn Sci 11:49–57. https://doi.org/10.1016/j.tics.2006.11.004.

Bzdok D, Heeger A, Langner R, Laird AR, Fox PT, Palomero-Gallagher N, Vogt BA, Zilles K, Eickhoff SB (2015): Subspecialization in the human posterior medial cortex. Neuroimage 106:55–71. http://dx.doi.Org/10.1016/j.neuroimage.2014.11.009.

Cavassila S, Deval S, Huegen C, van Ormondt D, Graveron-Demilly D (2001): Cramér-Rao bounds: an evaluation tool for quantitation. NMR Biomed 14:278–283. https://doi.org/10.1002/nbm.701.

Chan D, Gallaher LM, Moodley K, Minati L, Burgess N, Hartley T (2016): The 4 Mountains Test: A Short Test of Spatial Memory with High Sensitivity for the Diagnosis of Pre-dementia Alzheimer’s Disease. J Vis Exp 116:e54454. doi:10.3791/54454.

Dean HL (2006): Allocentric Spatial Referencing of Neuronal Activity in Macaque Posterior Cingulate Cortex. J Neurosci 26:1117–1127. https://doi.org/10.1523/JNEUROSCI.2497-05.2006.

Deichmann R, Gottfried J., Hutton C, Turner R (2003): Optimized EPI for fMRI studies of the orbitofrontal cortex. Neuroimage 19:430–441. https://doi.org/10.1016/S1053-8119(03)00073-9.

Diedenhofen B, Musch J (2015): cocor: A Comprehensive Solution for the Statistical Comparison of Correlations. Ed. Jake Olivier. PLoS One 10:e0121945. https://doi.org/10.1371/journal.pone.0121945.

Dienes Z, Mclatchie N (2018): Four reasons to prefer Bayesian analyses over significance testing. Psychon Bull Rev 25:207–218. https://doi.org/10.3758/s13423-017-1266-z.

Duarte JMN, Lei H, Mlynárik V, Gruetter R (2012): The neurochemical profile quantified by in vivo 1H NMR spectroscopy. Neuroimage 61:342–362. https://doi.org/10.1016/j.neuroimage.2011.12.038.

Duncan NW, Wiebking C, Munoz-Torres Z, Northoff G (2014): How to investigate neuro-biochemical relationships on a regional level in humans? Methodological considerations for combining functional with biochemical imaging. J Neurosci Methods 221:183–188. http://dx.doi.org/10.1016/j.jneumeth.2013.10.011.

Edden RAE, Puts NAJ, Harris AD, Barker PB, Evans CJ (2014): Gannet: A batch-processing tool for the quantitative analysis of gamma-aminobutyric acid-edited MR spectroscopy spectra. J Magn Reson Imaging 40:1445–1452. https://doi.org/10.1002/jmri.24478.

Eklund A, Nichols TE, Knutsson H (2016): Cluster failure: Why fMRI inferences for spatial extent have inflated false-positive rates. Proc Natl Acad Sci 113:7900–7905. https://doi.org/10.1073/pnas.1602413113.

Ekstrom AD, Huffman DJ, Starrett M (2017): Interacting networks of brain regions underlie human spatial navigation: a review and novel synthesis of the literature. J Neurophysiol 118:3328–3344. https://doi.org/10.1152/jn.00531.2017.

Enzi B, Duncan NW, Kaufmann J, Tempelmann C, Wiebking C, Northoff G (2012): Glutamate modulates resting state activity in the perigenual anterior cingulate cortex – A combined fMRI–MRS study. Neuroscience 227:102–109. http://dx.doi.org/10.1016/j.neuroscience.2012.09.039.

Epperson CN, Haga K, Mason GF, Sellers E, Gueorguieva R, Zhang W, Weiss E, Rothman DL, Krystal JH (2002): Cortical γ-Aminobutyric Acid Levels Across the Menstrual Cycle in Healthy Women and Those With Premenstrual Dysphoric Disorder. Arch Gen Psychiatry 59:851. doi:10.1001/archpsyc.59.9.851.

Esposito F, Aragri A, Latorre V, Popolizio T, Scarabino T, Cirillo S, Marciano E, Tedeschi G, Di Salle F (2009): Does the default-mode functional connectivity of the brain correlate with working-memory performances? Arch Ital Biol 147:11–20.

Farrant M, Nusser Z (2005): Variations on an inhibitory theme: phasic and tonic activation of GABAA receptors. Nat Rev Neurosci 6:215–229. https://doi.org/10.1038/nrn1625.

Fox KCR, Foster BL, Kucyi A, Daitch AL, Parvizi J (2018): Intracranial Electrophysiology of the Human Default Network. Trends Cogn Sci 22:307–324. https://doi.org/10.1016/j.tics.2018.02.002.

Fransson P, Marrelec G (2008): The precuneus/posterior cingulate cortex plays a pivotal role in the default mode network: Evidence from a partial correlation network analysis. Neuroimage 42:1178–1184. https://doi.org/10.1016/j.neuroimage.2008.05.059.

Futamura A, Honma M, Shiromaru A, Kuroda T, Masaoka Y, Midorikawa A, Miller MW, Kawamura M, Ono K (2018): Singular case of the driving instructor: Temporal and topographical disorientation. Neurol Clin Neurosci 6:16–18. https://doi.org/10.1111/ncn3.12166.

Greenhouse I, Noah S, Maddock RJ, Ivry RB (2016): Individual differences in GABA content are reliable but are not uniform across the human cortex. Neuroimage 139:1–7. http://dx.doi.org/10.1016/j.neuroimage.2016.06.007.

Hancu I (2009): Optimized glutamate detection at 3T. J Magn Reson Imaging 30:1155–1162. https://doi.org/10.1002/jmri.21936.

Hao X, Xu D, Bansal R, Dong Z, Liu J, Wang Z, Kangarlu A, Liu F, Duan Y, Shova S, Gerber AJ, Peterson BS (2013): Multimodal magnetic resonance imaging: The coordinated use of multiple, mutually informative probes to understand brain structure and function. Hum Brain Mapp 34:253–271. https://doi.org/10.1002/hbm.21440.

Harris AD, Puts NAJ, Edden RAE (2015a): Tissue correction for GABA-edited MRS: Considerations of voxel composition, tissue segmentation, and tissue relaxations. J Magn Reson Imaging 42:1431–1440. https://doi.org/10.1002/jmri.24903.

Harris AD, Saleh MG, Edden RAE (2017): Edited 1 H magnetic resonance spectroscopy in vivo: Methods and metabolites. Magn Reson Med 77:1377–1389. https://doi.org/10.1002/mrm.26619.

Harris AD, Puts NAJ, Anderson BA, Yantis S, Pekar JJ, Barker PB, Edden RAE (2015b): Multi-Regional Investigation of the Relationship between Functional MRI Blood Oxygenation Level Dependent (BOLD) Activation and GABA Concentration. Ed. Xi Luo. PLoS One 10:e0117531. https://doi.org/10.1371/journal.pone.0117531.

Hassabis D, Kumaran D, Maguire EA (2007): Using Imagination to Understand the Neural Basis of Episodic Memory. J Neurosci 27:14365–14374. https://doi.org/10.1523/JNEUROSCI.4549-07.2007.

Hassabis D, Maguire EA (2007): Deconstructing episodic memory with construction. Trends Cogn Sci 11:299–306. https://doi.org/10.1016/j.tics.2007.05.001.

van den Heuvel MP, Sporns O (2013): Network hubs in the human brain. Trends Cogn Sci 17:683–696. http://dx.doi.org/10.1016/j.tics.2013.09.012.

Hodgetts CJ, Shine JP, Lawrence AD, Downing PE, Graham KS (2016): Evidencing a place for the hippocampus within the core scene processing network. Hum Brain Mapp 37:3779–3794. https://doi.org/10.1002/hbm.23275.

Hodgetts CJ, Voets NL, Thomas AG, Clare S, Lawrence AD, Graham KS (2017): Ultra-High-Field fMRI Reveals a Role for the Subiculum in Scene Perceptual Discrimination. J Neurosci 37:3150–3159. https://doi.org/10.1523/JNEUROSCI.3225-16.2017.

Hodgetts CJ, Postans M, Shine JP, Jones DK, Lawrence AD, Graham KS (2015): Dissociable roles of the inferior longitudinal fasciculus and fornix in face and place perception. Elife 4:1–25. https://doi.org/10.7554/eLife.07902.001.

Hu Y, Chen X, Gu H, Yang Y (2013): Resting-State Glutamate and GABA Concentrations Predict Task-Induced Deactivation in the Default Mode Network. J Neurosci 33:18566–18573. https://doi.org/10.1523/JNEUROSCI.1973-13.2013.

Hutchison RM, Culham JC, Everling S, Flanagan JR, Gallivan JP (2014): Distinct and distributed functional connectivity patterns across cortex reflect the domain-specific constraints of object, face, scene, body, and tool category-selective modules in the ventral visual pathway. Neuroimage 96:216–236. http://dx.doi.org/10.1016/j.neuroimage.2014.03.068.

Ip IB, Berrington A, Hess AT, Parker AJ, Emir UE, Bridge H (2017): Combined fMRI-MRS acquires simultaneous glutamate and BOLD-fMRI signals in the human brain. Neuroimage 155:113–119. http://dx.doi.org/10.1016/j.neuroimage.2017.04.030.

Irish M, Halena S, Kamminga J, Tu S, Hornberger M, Hodges JR (2015): Scene construction impairments in Alzheimer’s disease – a unique role for the posterior cingulate cortex. Cortex 73:10–23. http://dx.doi.org/10.1016/j.cortex.2015.08.004.

Jagust WJ, Mormino EC (2011): Lifespan brain activity, β-amyloid, and Alzheimer’s disease. Trends Cogn Sci 15:520–526. https://doi.org/10.1016/j.tics.2011.09.004.

Jarosz AF, Wiley J (2014): What Are the Odds ? A Practical Guide to Computing and Reporting Bayes Factors. J Probl Solving 7:2–9. http://dx.doi.org/10.7771/1932-6246.1167.

Jenkinson M (2003): Fast, automated, N-dimensional phase-unwrapping algorithm. Magn Reson Med 49:193–197. https://doi.org/10.1002/mrm.10354.

Jenkinson M, Bannister P, Brady M, Smith S (2002): Improved Optimization for the Robust and Accurate Linear Registration and Motion Correction of Brain Images. Neuroimage 17:825–841. https://doi.org/10.1006/nimg.2002.1132.

Kravitz DJ, Saleem KS, Baker CI, Mishkin M (2011): A new neural framework for visuospatial processing. Nat Rev Neurosci 12:217–230. https://doi.org/10.1038/nrn3008.

Kriegeskorte N, Simmons WK, Bellgowan PSF, Baker CI (2009): Circular analysis in systems neuroscience: the dangers of double dipping. Nat Neurosci 12:535–540. https://doi.org/10.1038/nn.2303.

Landim RCG, Edden RAE, Foerster B, Li LM, Covolan RJM, Castellano G (2016): Investigation of NAA and NAAG dynamics underlying visual stimulation using MEGA-PRESS in a functional MRS experiment. Magn Reson Imaging 34:239–45. https://doi.org/10.1016/j.mri.2015.10.038.

Leech R, Sharp DJ (2014): The role of the posterior cingulate cortex in cognition and disease. Brain 137:12–32. https://doi.org/10.1093/brain/awt162.

Lipp I, Evans CJ, Lewis C, Murphy K, Wise RG, Caseras X (2015): The Relationship between Fearfulness, GABA+, and Fear-Related BOLD Responses in the Insula. Ed. Alexandra Kavushansky. PLoS One 10:e0120101. https://doi.org/10.1371/journal.pone.0120101.

Logothetis NK (2002): The neural basis of the blood-oxygen-level-dependent functional magnetic resonance imaging signal. Philos Trans R Soc B Biol Sci 357:1003–1037. https://doi.org/10.1098/rstb.2002.1114.

Marsman M, Wagenmakers EJ (2017): Bayesian benefits with JASP. Eur J Dev Psychol 14:545–555. https://doi.org/10.1080/17405629.2016.1259614.

McLean J, Krishnadas R, Batty GD, Burns H, Deans KA, Ford I, McConnachie A, McGinty A, McLean JS, Millar K, Sattar N, Shiels PG, Tannahill C, Velupillai YN, Packard CJ, Condon BR, Hadley DM, Cavanagh J (2012): Early life socioeconomic status, chronic physiological stress and hippocampal N-acetyl aspartate concentrations. Behav Brain Res 235:225–230. http://dx.doi.org/10.1016/j.bbr.2012.08.013.

Mescher M, Merkle H, Kirsch J, Garwood M, Gruetter R (1998): Simultaneous in vivo spectral editing and water suppression. NMR Biomed 11:266–72. https://doi.org/10.1002/(SICI)1099-1492(199810)11:6%3C266::AID-NBM530%3E3.0.CO;2-J.

Moffet J, Ross B, Arun P, Madhavarao C, Namboodiri A (2007): N-Acetylaspartate in the CNS: From neurodiagnostics to neurobiology. Prog Neurobiol 81:89–131. https://doi.org/10.1016/j.pneurobio.2006.12.003.

Mullins PG, McGonigle DJ, O’Gorman RL, Puts NAJ, Vidyasagar R, Evans CJ, Edden RAE (2014): Current practice in the use of MEGA-PRESS spectroscopy for the detection of GABA. Neuroimage 86:43–52. http://dx.doi.org/10.1016/j.neuroimage.2012.12.004.

Murray EA, Wise SP, Graham KS (2017): Representational specializations of the hippocampus in phylogenetic perspective. Neurosci Lett:http://dx.doi.org/10.1016/j.neulet.2017.04.065. https://doi.org/10.1016/j.neulet.2017.04.065.

Nakagawa S (2004): A farewell to Bonferroni: the problems of low statistical power and publication bias. Behav Ecol 15:1044–1045. https://doi.org/10.1093/beheco/arh107.

Palombo DJ, Hayes SM, Peterson KM, Keane MM, Verfaellie M (2018): Medial Temporal Lobe Contributions to Episodic Future Thinking: Scene Construction or Future Projection? Cereb Cortex 28:447–458. https://doi.org/10.1093/cercor/bhw381.

Parvizi J, Van Hoesen GW, Buckwalter J, Damasio A (2006): Neural connections of the posteromedial cortex in the macaque. Proc Natl Acad Sci 103:1563–1568. https://doi.org/10.1073/pnas.0507729103.

Passarelli L, Rosa MGP, Bakola S, Gamberini M, Worthy KH, Fattori P, Galletti C (2018): Uniformity and Diversity of Cortical Projections to Precuneate Areas in the Macaque Monkey: What Defines Area PGm? Cereb Cortex 28:1700–1717. https://doi.org/10.1093/cercor/bhx067.

Patel TB, Clark JB (1979): Synthesis of N-acetyl-l-aspartate by rat brain mitochondria and its involvement in mitochondrial/cytosolic carbon transport. Biochem J 184:539–546.

Rae CD (2014): A Guide to the Metabolic Pathways and Function of Metabolites Observed in Human Brain 1H Magnetic Resonance Spectra. Neurochem Res 39:1–36. https://doi.org/10.1007/s11064-013-1199-5.

Raichle ME (2015): The Brain’s Default Mode Network. Annu Rev Neurosci 38:433–447. https://doi.org/10.1146/annurev-neuro-071013-014030.

Raichle ME, MacLeod AM, Snyder AZ, Powers WJ, Gusnard DA, Shulman GL (2001): A default mode of brain function. Proc Natl Acad Sci USA 98:676–682. https://doi.org/10.1073/pnas.98.2.676.

Ranganath C, Ritchey M (2012): Two cortical systems for memory-guided behaviour. Nat Rev Neurosci 13:713–726. https://doi.org/10.1038/nrn3338.

Sato N, Sakata H, Tanaka YL, Taira M (2010): Context-Dependent Place-Selective Responses of the Neurons in the Medial Parietal Region of Macaque Monkeys. Cereb Cortex 20:846–858. https://doi.org/10.1093/cercor/bhp147.

Schacter DL, Addis DR, Hassabis D, Martin VC, Spreng RN, Szpunar KK (2012): The Future of Memory: Remembering, Imagining, and the Brain. Neuron 76:677–694. https://doi.org/10.1016/j.neuron.2012.11.001.

Shine JP, Hodgetts CJ, Postans M, Lawrence AD, Graham KS (2015): APOE-ε4 selectively modulates posteromedial cortex activity during scene perception and short-term memory in young healthy adults. Sci Rep 5:16322. https://doi.org/10.1038/srep16322.

Simmons ML, Frondoza CG, Coyle JT (1991): Immunocytochemical localization of N-acetyl-aspartate with monoclonal antibodies. Neuroscience 45:37–45.

Smith SM (2002): Fast robust automated brain extraction. Hum Brain Mapp 17:143–155. https://doi.org/10.1002/hbm.10062.

Spreng RN, Sepulcre J, Turner GR, Stevens WD, Schacter DL (2013): Intrinsic Architecture Underlying the Relations among the Default, Dorsal Attention, and Frontoparietal Control Networks of the Human Brain. J Cogn Neurosci 25:74–86. https://doi.org/10.1162/jocn_a_00281.

Spreng RN (2012): The Fallacy of a “Task-Negative” Network. Front Psychol 3:145. https://doi.org/10.3389/fpsyg.2012.00145.

Spreng RN, Mar RA, Kim ASN (2009): The Common Neural Basis of Autobiographical Memory, Prospection, Navigation, Theory of Mind, and the Default Mode: A Quantitative Meta-analysis. J Cogn Neurosci 21:489–510. https://doi.org/10.1162/jocn.2008.21029.

Stagg CJ, Bachtiar V, Johansen-Berg H (2011): The Role of GABA in Human Motor Learning. Curr Biol 21:480–484. https://doi.org/10.1016/j.cub.2011.01.069.

Stagg CJ, Rothman DL eds. (2014): Magnetic Resonance Spectroscopy: Tools for Neurosceicne Research and Emerging Clinical Applications. Elsevier.

Suzuki K, Yamadori A, Hayakawa Y, Fujii T (1998): Pure topographical disorientation related to dysfunction of the viewpoint dependent visual system. Cortex 34:589–599. https://doi.org/10.1016/S0010-9452(08)70516-1.

Terpstra M, Cheong I, Lyu T, Deelchand DK, Emir UE, Bednařík P, Eberly LE, Öz G (2016): Test-retest reproducibility of neurochemical profiles with short-echo, single-voxel MR spectroscopy at 3T and 7T. Magn Reson Med 76:1083–1091. https://doi.org/10.1002/mrm.26022.

Tkáč I, Öz G, Adriany G, Uğurbil K, Gruetter R (2009): In vivo 1 H NMR spectroscopy of the human brain at high magnetic fields: Metabolite quantification at 4T vs. 7T. Magn Reson Med 62:868–879. https://doi.org/10.1002/mrm.22086.

Utevsky A V, Smith D V, Huettel S a (2014): Precuneus Is a Functional Core of the Default-Mode Network. J Neurosci 34:932–940. https://doi.org/10.1523/JNEUROSCI.4227-13.2014.

Wetzels R, Wagenmakers E-J (2012): A default Bayesian hypothesis test for correlations and partial correlations. Psychon Bull Rev 19:1057–1064. https://doi.org/10.3758/s13423-012-0295-x.

Wilson M, Reynolds G, Kauppinen R a, Arvanitis TN, Peet AC (2011): A constrained least-squares approach to the automated quantitation of in vivo 1 H magnetic resonance spectroscopy data. Magn Reson Med 65:1–12. https://doi.org/10.1002/mrm.22579.

Woolrich MW, Behrens TEJ, Beckmann CF, Jenkinson M, Smith SM (2004): Multilevel linear modelling for FMRI group analysis using Bayesian inference. Neuroimage 21:1732–1747. https://doi.org/10.1016/j.neuroimage.2003.12.023.

Zhang Y, Brady M, Smith S (2001): Segmentation of brain MR images through a hidden Markov random field model and the expectation-maximization algorithm. IEEE Trans Med Imaging 20:45–57. doi:10.1109/42.906424.

